# GLASS: A Graph Learning Algorithm for Screening Splice-Aware Alignments of Long-Read RNA-seq data

**DOI:** 10.1101/2025.04.07.647681

**Authors:** Jiahao Li, Guojun Li, Ting Yu

**Author notes:** Contributing authors.

## Abstract

With the continuous development of third-generation RNA-seq, obtaining accurate splice-aware alignments for the long RNA-seq reads to reference genomes has become one of the major challenges in transcriptomic analysis. To mitigate the erroneous alignments arising from existing long-read RNA-seq aligners, we propose GLASS, a novel splice-aware alignments filtering approach based on third-generation transcriptome data. GLASS introduces a newly designed ‘Read-AS Map’ model and integrates graph machine learning techniques for detecting and removing falsely spliced aligned reads from alignment files. Experimental results demonstrate that GLASS significantly reduces the error rate of spliced alignmnt and enhance the accuracy of subsequent transcriptome reconstruction. Additionally, GLASS demonstrates strong generalization ability in data from species such as Mus musculus and Arabidopsis thaliana, providing a new approach for the efficient processing of third-generation transcriptome data and offering more reliable data support for subsequent gene expression analysis and functional studies.

## 1 Introduction

In recent years, long-read sequencing technologies have garnered widespread attention and have achieved significant applications across various fields [1, 2]. With the ongoing advancement of transcriptomics research, both second- and third-generation RNA sequencing technologies (RNA-Seq) [3] have become essential tools for studying gene expression and many different aspects of RNA biology [4]. Second-generation RNA-Seq technologies, such as the Illumina platform, are widely used in transcriptomic data analysis due to their high throughput, low cost, and well-established analytical workflows [5]. However, despite the remarkable achievements of second-generation technologies in transcriptomics, third-generation RNA-Seq (such as PacBio and Oxford Nanopore technologies) [6, 7] is gradually demonstrating distinct advantages. Third-generation RNA-Seq allows for the direct reading of full-length transcripts, reducing the reliance on assembly and offering unparalleled advantages in capturing complex transcripts and novel isoforms [8].

In transcriptomic analysis based on reference genomes, alignment of transcripts to the reference genome is a crucial step. It directly impacts the quantification of gene expression, the identification of transcripts, and the analysis of splicing. Commonly used second-generation RNA-seq aligners include HISAT2 [9] and STAR [10], while the most popular third-generation tool is Minimap2 [11]. However, these RNA-seq aligners are not flawless. Due to inherent differences between an individual’s genome and the reference genome, as well as inevitable errors during sequencing and alignment algorithmic limitations, it is impossible to perfectly align raw RNA-seq data to the existing reference genome.

As a result, the alignment files generated during RNA-seq data analysis often contain alignments that are inconsistent with the actual biological characteristics or reference annotations. These results are typically considered “incorrect alignments”, as they not only fail to reflect the true biological situation but may also affect the accuracy of transcriptome reconstruction. Such erroneous alignments act as noise in the alignment files, potentially masking valuable biological information or even leading the analysis to incorrect conclusions. Therefore, accurately identifying and removing these incorrect alignments-especially those that are inconsistent with the reference annotations-has become an important challenge for improving data quality and downstream analysis precision in RNA-seq studies.

Alternative splicing (AS) is a mechanism through which exons from a single pre-mRNA are differentially combined to produce multiple mRNAs, thereby enhancing proteome diversity and regulating gene expression in higher eukaryotes [12]. In this research, the term “alternative splicing event” is occasionally used to refer to specific instances of AS. We define non-reference AS events as those absent in reference annotations, while reference AS events are those documented in reference annotations.

To quantify the defect of RNA-seq aligners, we conducted statistical analysis on data from the same species and tissue type. We found that for the second-generation RNA-seq data SRR4235527, after processing with HISAT2, the non-reference AS rate was approximately 6.24%. In contrast, for third-generation data NA12878, processed with Minimap2, the rate was about 10.68%. This result suggests that third-generation data have a higher proportion of suspicious alignments, leading to more spurious AS events, and also indicates that third-generation RNA-seq still have significant room for improvement in this area. The operational commands for HISAT2 and Minimap2 are provided in the supplementary materials, while the versions of the reference materials are listed in **Table 2**.

**Table 1.**
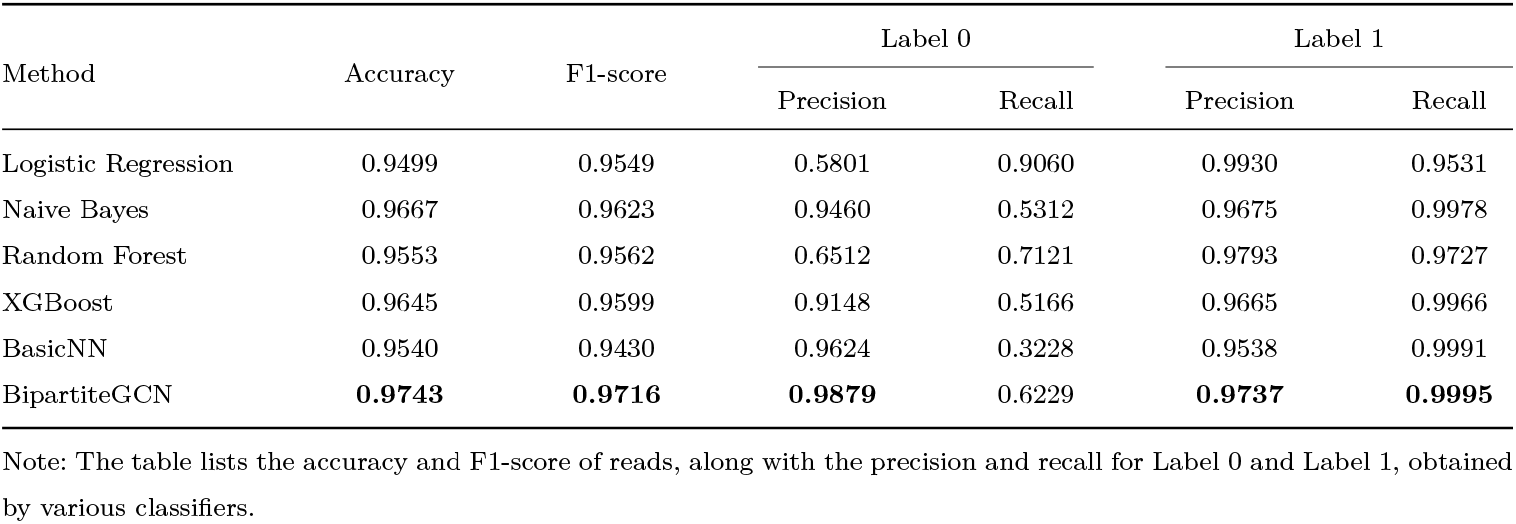
Comparison of results of different classifiers on NA12878.

| Method | Accuracy | F1-score | Label 0 |  | Label 1 |  |
| --- | --- | --- | --- | --- | --- | --- |
|  |  |  | Precision | Recall | Precision | Recall |
| Logistic Regression | 0.9499 | 0.9549 | 0.5801 | 0.9060 | 0.9930 | 0.9531 |
| Naive Bayes | 0.9667 | 0.9623 | 0.9460 | 0.5312 | 0.9675 | 0.9978 |
| Random Forest | 0.9553 | 0.9562 | 0.6512 | 0.7121 | 0.9793 | 0.9727 |
| XGBoost | 0.9645 | 0.9599 | 0.9148 | 0.5166 | 0.9665 | 0.9966 |
| BasicNN | 0.9540 | 0.9430 | 0.9624 | 0.3228 | 0.9538 | 0.9991 |
| BipartiteGCN | **0.9743** | **0.9716** | **0.9879** | 0.6229 | **0.9737** | **0.9995** |

Note: The table lists the accuracy and F1-score of reads, along with the precision and recall for Label 0 and Label 1, obtained by various classifiers.

**Table 2.** Summary of Experimental Materials and Data.

| Homo sapiens |  |  |  |
| --- | --- | --- | --- |
| Train Data ID | Test Data ID | Validation Data ID | Reference ID |
| SRR25180684 | NA12878 |  |  |
| ERR4706159 | ERR6053080 |  |  |
| SRR23347361 | SRR30786404 | SRR1163657 | GCF_000001405.39 |
| DRR228524 | SRR30692516 |  |  |
| ERR9286493 | SRR20637410 |  |  |
| Mus musculus |  | Arabidopsis thaliana |  |
| Test Data ID | Reference ID | Test Data ID | Reference ID |
| SRR26545106 |  | SRR28341944 | GCF_000001735.4 |
| SRR17253008 | GCA_000001635.9 |  |  |
| SRR30692520 |  | DRR424742 |  |

For data generated by second-generation RNA-seq aligners, previous works, such as EASTR [13], have proposed solutions to address this problem metioned above. EASTR first classifies sequences based on the sequence similarity of upstream and downstream flanking regions. It then evaluates the spliced alignments by considering the frequency of sequence occurrences in the reference genome, thereby enabling the identification and removal of potential spurious spliced alignments.

However, third-generation RNA-Seq also face the simliar challenge as above mentioned statistical results. Notably, EASTR is better suited for second-generation RNA-Seq, and its core algorithm has significant limitations when applied to third-generation RNA-Seq data. Analysis of EASTR revealed that its good performance with second-generation RNA-Seq data can be attributed the short reads of second-generation sequencing, which are more prone to alignment errors when encountering repetitive regions. In contrast, third-generation RNA-Seq data features long that are better able to retain complete transcript information and can span across multiple repetitive regions. As a result, EASTR is less effective at detecting and removing splicing errors in third-generation data. This explains why EASTR performs well with second-generation data, but its effectiveness is diminished when applied to third-generation data. Consequently, designing tools specifically adapted to the unique characteristics of third-generation transcriptomics has become an important focus of current transcriptomic research.

From another perspective, for tools like SpliceAI [14], Splam [15], and DeltaSplice [16], they typically rely solely on sequence information for analysis, without considering the global context of the data. Additionally, most of these tools focus primarily on classifying the accuracy of splice sites, without delving into how to integrate information from multiple AS events within a read. The effectiveness of using read classification methods to find low-quality alignments has been a topic of ongoing debate. This study aims to address this issue by exploring how effective conducting read classification strategies to filter low-quality alignments.

In terms of data properties, second-generation sequencing and third-generation sequencing) data also exhibit significant differences. We found that among all reads that support AS, the proportion of reads spanning multiple splice variants in third-generation RNA-Seq data (NA12878) was 84.47%, whereas the same proportion in second-generation RNA-Seq data (SRR4235527) was only 4.09%. Leveraging this unique characteristics of third-generation sequencing data, we introduced the ‘Read-AS Map’ model, drawing inspiration from graph classification methods, which have been widely applied in various fields, such as predicting molecular properties of compounds [17, 18].

In summary, to address the aforementioned challenges, we proposed GLASS, a novel third-generation transcriptome filtering method that integrates graph learning algorithms. GLASS takes advantages of the ability of long reads to span multiple AS events, ensuring the effective utilization of the long-read characteristics inherent in third-generation transcriptome data.

## 2 Results

### 2.1 Graph-Driven BipartiteGCN Boosts Read Label Prediction

As is well known, chromosomal information is highly specific to individual species. Our goal, however, is to enable the model to learn more generalized patterns, rather than being restricted to species-specific genomic features. Therefore, directly embedding chromosomal information as a feature into the model may be unreasonable, as it could hinder the model’s ability to generalize across different species. This presents the first challenge in the design of our algorithm: how to retain the relative positions of reads and AS events as much as possible.

Graphs are an excellent model for representing complex relationships and structural information. To address the issue mentioned above, we designed a bipartite graph model ‘Read-AS map’. In this model, reads and AS events are treated as nodes, and edges are created between them, thereby preserving their positional information without relying on chromosomal structure. This approach enables the model to capture the relationships between reads and AS events while avoiding the species-specific limitations imposed by direct reliance on chromosomal data. In practice, the choice of the graph model has proven successful. We could demonstrate the superiority of ‘Read-AS Map’ through two aspects.

On the one hand, we compared our method with other classifiers, including Logistic Regression [19], Naive Bayes[20], Random Forest[21], XGBoost [22], and BasicNN (a neural network classifier composed of regular linear layers) [23].

The experimental results presented in **Tabel 1** demonstrate the superior performance of the BipartiteGCN model across multiple evaluation metrics. Notably, BipartiteGCN achieves the highest accuracy (0.9743) and F1-score (0.9716) among all classifiers, outperforming traditional machine learning methods that we used in the experiments. Particularly impressive is BipartiteGCN’s performance in read with label 0 prediction, where it achieves a remarkable precision of 0.9879 while maintaining a recall of 0.6229, striking an optimal balance that other models fail to achieve. For read with label 1, BipartiteGCN maintains near-perfect recall (0.9995) while preserving high precision (0.9737). The consistent superiority of BipartiteGCN across both classes highlight ability of graph model to effectively capture complex relationships in the data while minimizing false positives, which laid the foundation for us to conduct further experiments and analysis. The design of the label is detailed in **Fig. 4**, and we have included the formulas for accuracy, F1-score, precision and recall in the supplementary materials.

On the other hand, our further experiments of have validated the model’s transferability across different species, confirming that our design concept is both scientifically sound and efficient. The relevant experimental results were detailed in **Fig. 1** and **Fig. 2**.

**Fig. 1.**
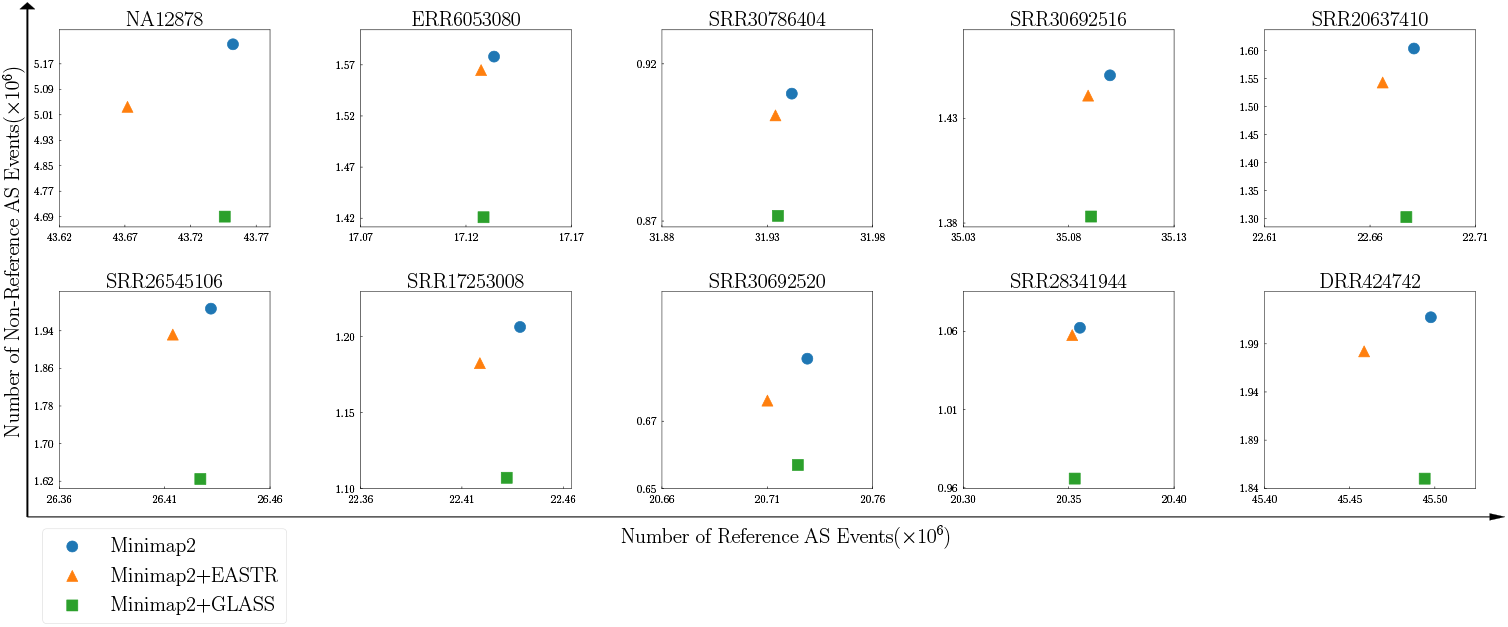
Impact of GLASS on the number of reference and non-reference AS events from alignment files. The horizontal axis represents the number of reference AS events, and the vertical axis represents the number of non reference AS events.

### 2.2 GLASS Effectively Removed Non-reference AS

To comprehensively demonstrate the advantages and capabilities of GLASS in filtering read with low-quality alignment of third-generation RNA-seq data, we conducted a detailed comparison with EASTR, a widely used tool for filtering alignment files. To ensure the results are broadly applicable and representative, we selected multiple real RNA-seq datasets, covering different species and experimental platforms, for testing. In the benchmark, we mainly compare the performance of GLASS with EASTR through two main experiments.

To effectively evaluate the performance of GLASS and EASTR in removing non-reference AS, we conducted a statistic of results before and after filtering of these two tools. Our focus was on whether they could accurately identify and filter out AS that do not align with the reference annotations while retaining the reference-consistent AS-those that represent true biological signals. Through this process, we aim to highlight the advantages of GLASS in enhancing the accuracy of alignment results.

Specifically, we first identify all reads that undergo AS process, then filter for primary alignments and those that can distinguish the ‘+’ and ‘-’ strands. Next, we extract all the AS from these reads and compare them with the reference annotations in the GTF file. Through this process, all AS events are classified as reference or non-reference, thereby establishing the foundation for subsequent statistical analyses.

We compared three workflows (Minimap2, Minimap2+EASTR, and Minimap2+GLASS) using ten sets of sample data, and we calculated the indicators defined in the previous section, as shown in **Fig. 1**. Initially, in the NA12878 sample, Minimap2 detected 43.75 million reference AS events and approximately 5.23 million non-reference AS events. When using Minimap2+GLASS, we observed significant improvements, particularly in terms of retaining reference AS and reducing non-reference AS. Specifically, Minimap2+GLASS detected 43.74 million reference AS events, more than the 43.67 million detected by Minimap2+EASTR, thus preserving more valid RNA-seq information. Furthermore, compared to the number of non-reference AS events identified by Minimap2, the number decreased by 3.8% to 5.03 million with Minimap2+EASTR. In contrast, Minimap2+GLASS exhibited better performance, with the number of non-reference AS events reduced by 10.1%, reaching 4.69 million.

This optimization trend was consistent across other species as well; for example, in the mouse sample SRR26545106, the number of non-reference AS events was reduced by 18.2% compared to Minimap2, and in the Arabidopsis sample DRR424742, Minimap2+GLASS showed an 8.3% reduction in the number of non-reference AS events. These findings indicate that Minimap2+GLASS, through its unique algorithmic design, significantly reduces the proportion of non-reference AS while maintaining the sensitivity of the original RNA-seq aligners thereby providing more accurate splice site information for subsequent transcript reconstruction.

### 2.3 GLASS Improved the results of Transcriptome Assembly

The quality of alignment file preprocessing is crucial for ensuring accurate RNA-seq transcript assembly. High-quality alignment files significantly reduce false-positive AS events caused by sequencing errors or individual genome biases, thereby improving the reliability of downstream transcript reconstruction [24].

Precise splice junction anchoring - the unambiguous alignment of reads spanning exon-intron boundaries - serves as a prerequisite for accurate AS identification, while alignments with excessive mismatch rates may introduce spurious exon-intron boundary signals [25]. Notably, although modern assembly algorithms incorporate stringent filtering mechanisms, their effectiveness remains highly dependent on input alignment quality. StringTie2 is a widely used transcript assembly tool in RNA-seq data analysis, known for its efficient assembly algorithm that made it well-suited for complex transcriptomes. As demonstrated in StringTie2, this state-of-the-art assembler employs a dual strategy to actively eliminate questionable spliced alignments: (1) imposing elevated coverage thresholds (25% higher than the default 1 read per bp) for spliced alignments with >1% mismatch rates (substantially exceeding Illumina’s 0.5% error rate), and (2) requiring extended anchor sequences (25 bp vs. default 10 bp) for alignments spanning ultra-long introns (>100, 000bp). However, when processing RNA-seq data, errors in the alignment files or limitations of the assembly algorithm may still result in inaccuracies in the assembled transcripts generated by StringTie2. This proactive filtering mechanism reveals that residual low-quality signals in raw alignment files may still interfere with model performance, even within high-stringency algorithmic frameworks. Therefore, developing precise filtering strategies to remove incorrect alignments, thereby improving the completeness and accuracy of the assembly, is crucial for enhancing the quality of RNA-seq analysis results.

In this study, we applied two data filtering tools, GLASS and EASTR, to the alignment files, followed by transcriptome assembly using StringTie2 [26], and evaluated their impact on assembly quality using gffcompare [27]. The versions of the reference materials we used in the evaluation process are listed in **Table 2**. We specifically focused on several key metrics, including the proportion of non-reference exons, the proportion of non-reference introns, and the precision and recall of matching intron chains. It is noteworthy that the matching intron chain is one of the most important metrics in the evaluation of transcriptome reconstruction algorithms, with numerous transcriptome assembly studies using this metric as the primary evaluation criterion. By analyzing these metrics in detail, we could assess the effect of the filtering tools on the assembly outcomes and ensure that, after filtering, the alignment files retain as much biological signal as possible, minimizing the loss of critical information.

The evaluation results based on several RNA-seq datasets demonstrate that StringTie2 with GLASS-cleaned alignments exhibits significant advantages. As shown in Figure 2, running StringTie2 with the alignment file cleaned by GLASS exhibited a significantly lower error rate. Notably, in the NA12878 sample, the non-reference exon rate of StringTie2 with GLASS-cleaned alignments was 24.3%, representing a 14.4% reduction compared to StringTie2 with raw alignments (27.8%). In contrast, StringTie2 with EASTR-cleaned alignments (26.8%) achieved only a 3.6% reduction. Similarly, the non-reference intron rate (4.7%) was significantly lower than that of StringTie2 with raw alignments (7.2%) and StringTie2 with EASTR-cleaned alignments (6.6%).

**Fig. 2.**
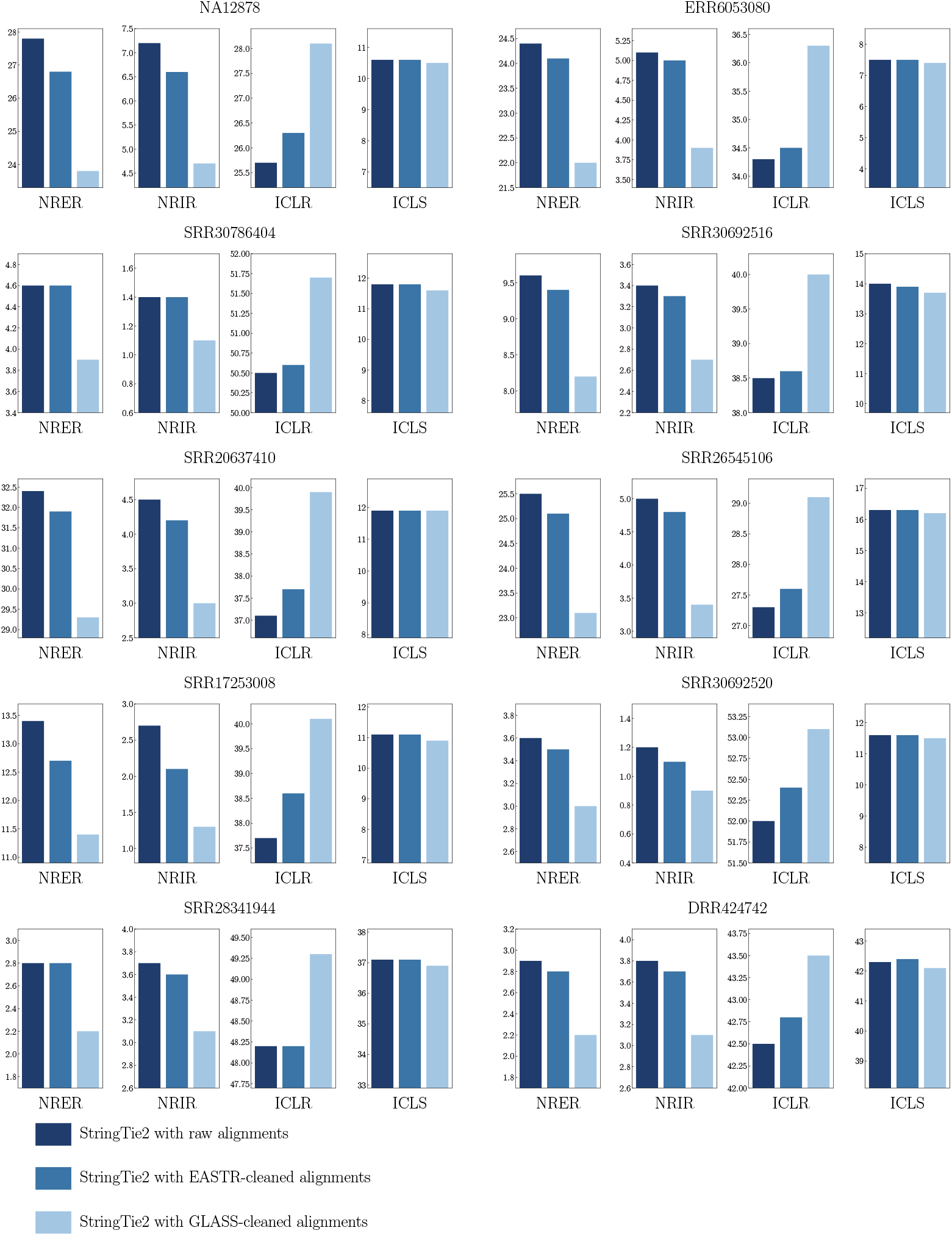
Influence of filtering on the assembling results before and after. As shown in the figure, NRER represents non reference exons rate, NRIR stands for non reference introns rate, ICLR stands for intron chain level precision and ICLS stands for intron chain level sensitivity. Please note that the data in this figure are all presented as percentages.

At the level of matching intron chains, StringTie2 with GLASS-cleaned alignments achieved higher accuracy. The intron chain accuracy for the NA12878 sample reached 28.1%, surpassing StringTie2 with raw alignment (25.7%) by a 9.3% improvement, while StringTie2 with EASTR-cleaned alignments only improved to 26.3%.

In more detail, across the five human datasets used in the experiment, StringTie2 with GLASS-cleaned alignments achieved an average reduction of 15.9% in the non-reference exon rate and a 33.4% average reduction in the non-reference intron rate compared to the StringTie2 with raw alignment. The improvement in the precision of matching intron chains was also significant, with StringTie2 with GLASS-cleaned achieving an average increase of 6.3% in precision across all human samples, while maintaining comparable sensitivity to StringTie2 with raw alignment and with EASTR-cleaned methods (sensitivity difference < 0.3%).

Moreover, this advantageous pattern is consistently reinforced across other non-human datasets, further validating the method’s ability to significantly enhance multi-level recognition accuracy while preserving detection sensitivity.

### 2.4 BipartiteGCN is a reliable predictive model

To further assess the stability and reliability of the model’s performance, we conducted an additional experiment aimed at demonstrating its scientific validity. As shown in **Fig. 3**, we introduced a separate validation set during the training process to monitor the model’s performance on unseen data in real time. In addition to optimizing the model using the training set, we plotted the loss curve for the validation set, which allowed us to directly observe the model’s generalization ability and verify the absence of overfitting. Specifically, as depicted in the loss curve, the validation loss steadily decreased throughout training and eventually stabilized, indicating that the model generalizes effectively without overfitting.

**Fig. 3.**
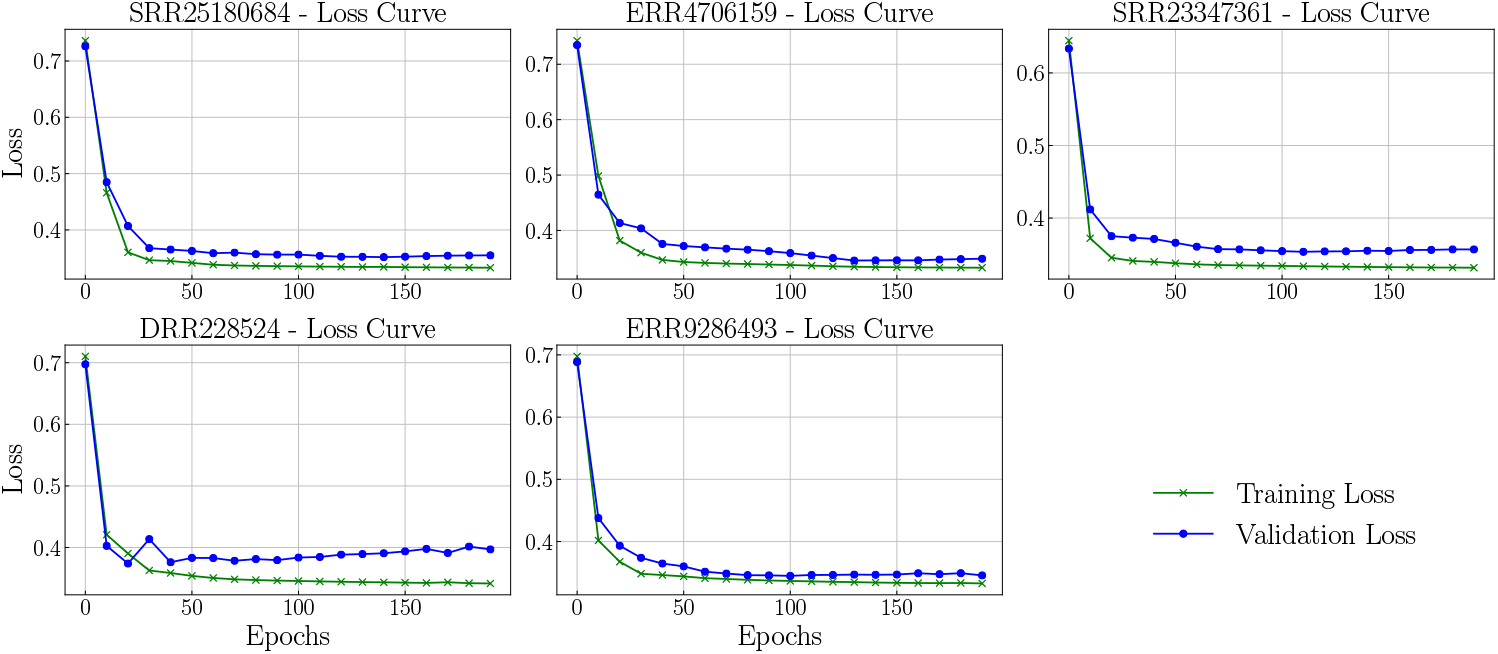
Loss curves of five training datasets. The horizontal axis represents the number of epochs, and the vertical axis represents the value of loss.

## 3 Methods

GLASS processes an alignment file (in BAM format) generated by splice-aware RNA-seq aligners, such as Minimap2, to identify and remove the potentionally erroneous spliced alignments, procuding a clean BAM file. In this study, we focused exclusively on reads within the alignment file satisfing three specific criteria: (1) spanning AS events, (2) meeting the requirements of primary alignments, and (3) having the characteristics that can be distinguished between the ‘+’ and ‘-’ strands. All reads that met these characteristics would be further analyzed.

### 3.1 Label and Feature Design

A positive sample refers to a read where the AS matches completely or partially with the AS in annotation, while a negative sample refers to a read that does not match any of the AS events in annotation. The following **Fig. 4** shows how labels are designed.

**Fig. 4.**
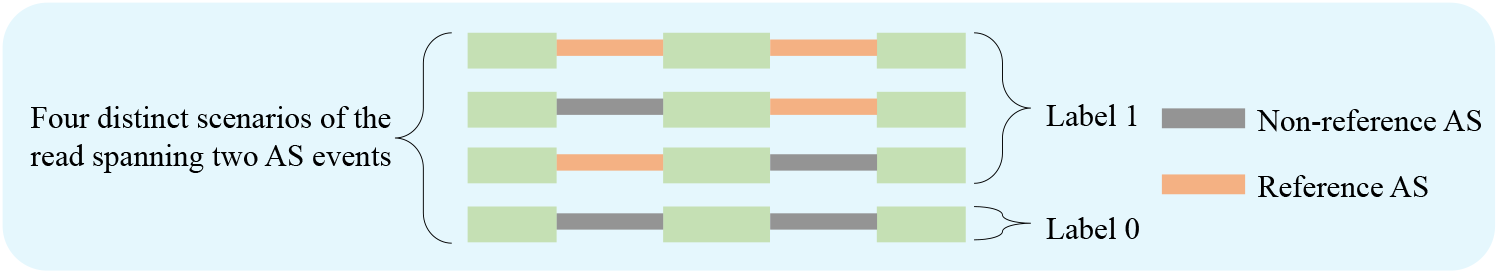
Schematic diagram of label design. The figure illustrates the reads spanning two AS events as examples. These two AS events correspond to four potential scenarios, through which the diagram visually demonstrates the design logic of the labeling strategy.

Before introducing the specific methods, we will provide a description of our research route. First, we will introduce the principles of label design, ensuring that the label system is efficient, aligning seamlessly with the task requirements. Next, we will explore how to construct the graph structure tailored to the task, allowing for effective representation of relationships between vertexs of different types. Following this, we will outline the specific computational steps, covering the process from data input to result prediction. Subsequently, we will describe the core components of the model architecture. Finally, we will summarize the evaluation strategies to ensure the model’s performance meets the expected objectives.

The features are designed based on the information from the alignment file generated by Minimap2, and the alternative splicing. The detailed feature design scheme is provided in the supplementary materials.

### 3.2 Construction of ‘Read-AS Map’

The ‘Read-AS Map’ is a bipartite graph *G* = (*X, Y, E*), which is constructed from a alignment file:

- *X* = *{x*_1_, *x*_2_, …, *x*_*n*_*}* is the set of reads, with each node *x*_*i*_ corresponding to a read (e.g. the corresponding primary alignment) in the alignment file.
- *Y* = *{y*_1_, *y*_2_, …, *y*_*m*_*}* is the set of AS events that identified by Minimap2, with each node *y*_*i*_ corresponding to an AS event in the alignment file.
- *E* is the set of directed edges of the graph. Based on the results from alignment file, if *y*_*j*_ is an AS event of *x*_*i*_, then mutually reaching directed edges will be constructed between *x*_*i*_ and *y*_*j*_.

The image below **Fig. 5**: (C), provides a schematic representation of the graph defined above.

**Fig. 5.**
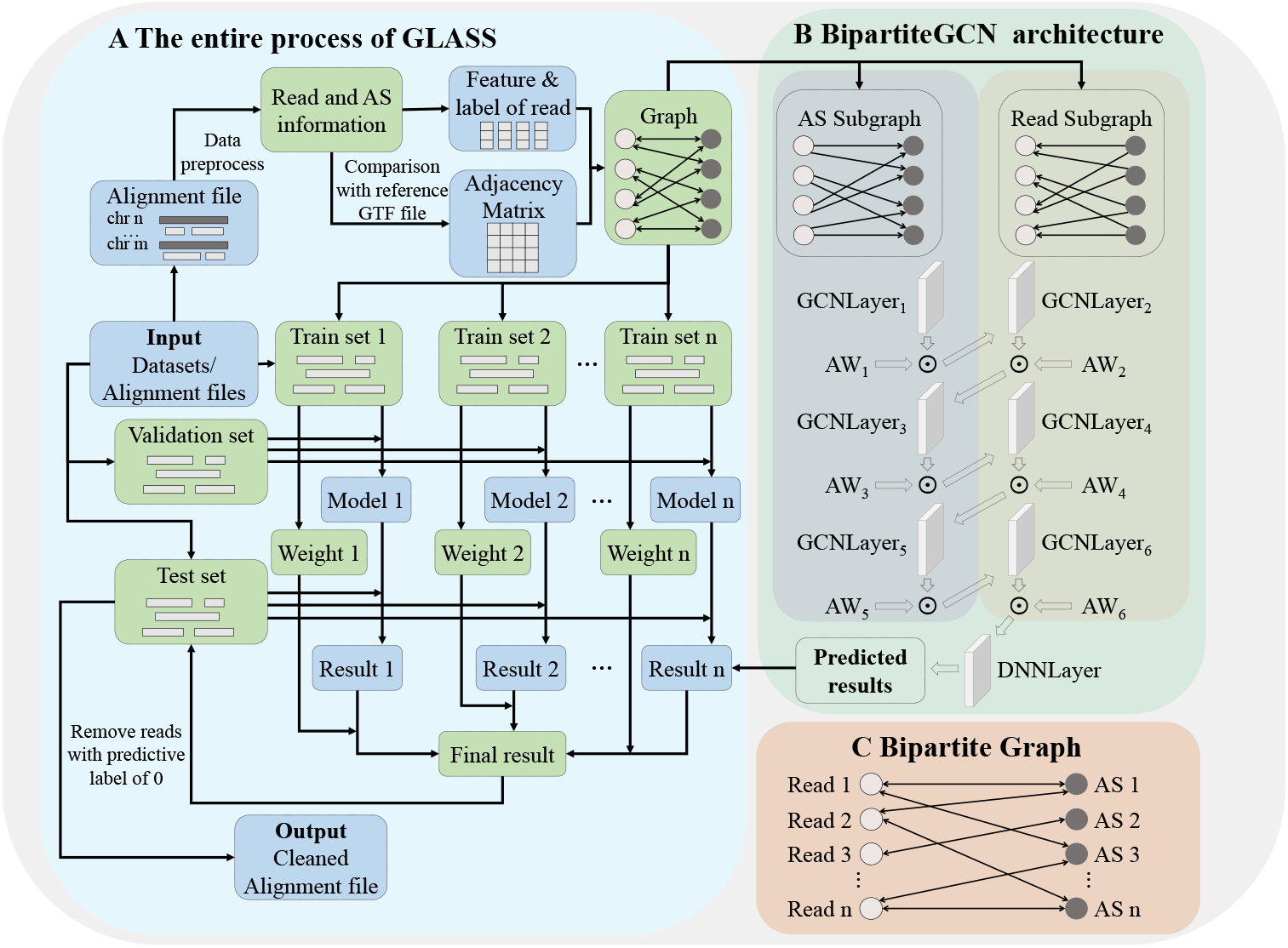
Full process of GLASS and framework of BipartiteGCN. This figure consists of three core components: the entire process of GLASS **(A)**, which describes the full processing steps from raw data input, model prediction to the final output; the BipartiteGCN architecture **(B)**, which details the hierarchical structure of the BipartiteGCN, including the input layer, graph convolution layers, and the integration of the attention mechanism (AW_i_ represents the attention weight of GCNLayer_i_); and the bipartite graph structure **(C)**, which highlights the model’s key bidirectional interaction mechanism, constructing a bipartite graph using two types of heterogeneous nodes (read and AS) to support cross-type node feature propagation.

### 3.3 Overview of the computational procedure

The operational steps are summarized below, with **Fig. 5**: (A) and **Fig. 5**: (B) providing a schematic representation of the approach framework.

- Model input: The model input consists of the features for each node *x* in *X*, and an adjacency matrix *A*[*i, j*] representing the relationships between nodes *x*_*i*_ and *y*_*j*_ in the bipartite graph.
- Graph Convolution Operation: The node feature representations are updated through multiple layers of graph convolution [28].
- Attention Mechanism: By learning attention weights at different layers, the algorithm is able to assign importance to each layer’s contribution [29].
- Label Smoothing Loss: Since the number of samples with a label of zero is relatively small in the dataset, we applied label smoothing regularization [30] to reduce overfitting.
- Ensemble Learning: The final prediction is obtained by performing a weighted average of the results from multiple models.
- Alignment File Cleaning: By integrating the prediction of read labels, the algorithm performs filtering on the alignment file, generating the final output.

### 3.4 BipartiteGCN Model

This model utilizes a bipartite graph structure, where node features are updated through bidirectional propagation via two types of edges:

- Forward Convolution (from “read” nodes to “AS” nodes).
- Backward Convolution (from “AS” nodes to “read” nodes).

Initially, node features are updated layer by layer using a Graph Convolutional Network (GCN), where each layer aggregates the features of the previous layer through normalization of the adjacency matrix, weighted combinations, and applies a learned weight matrix for linear transformation, followed by a nonlinear transformation using the ReLU activation function.

#### Algorithm 1 Bipartite Graph GCN Model

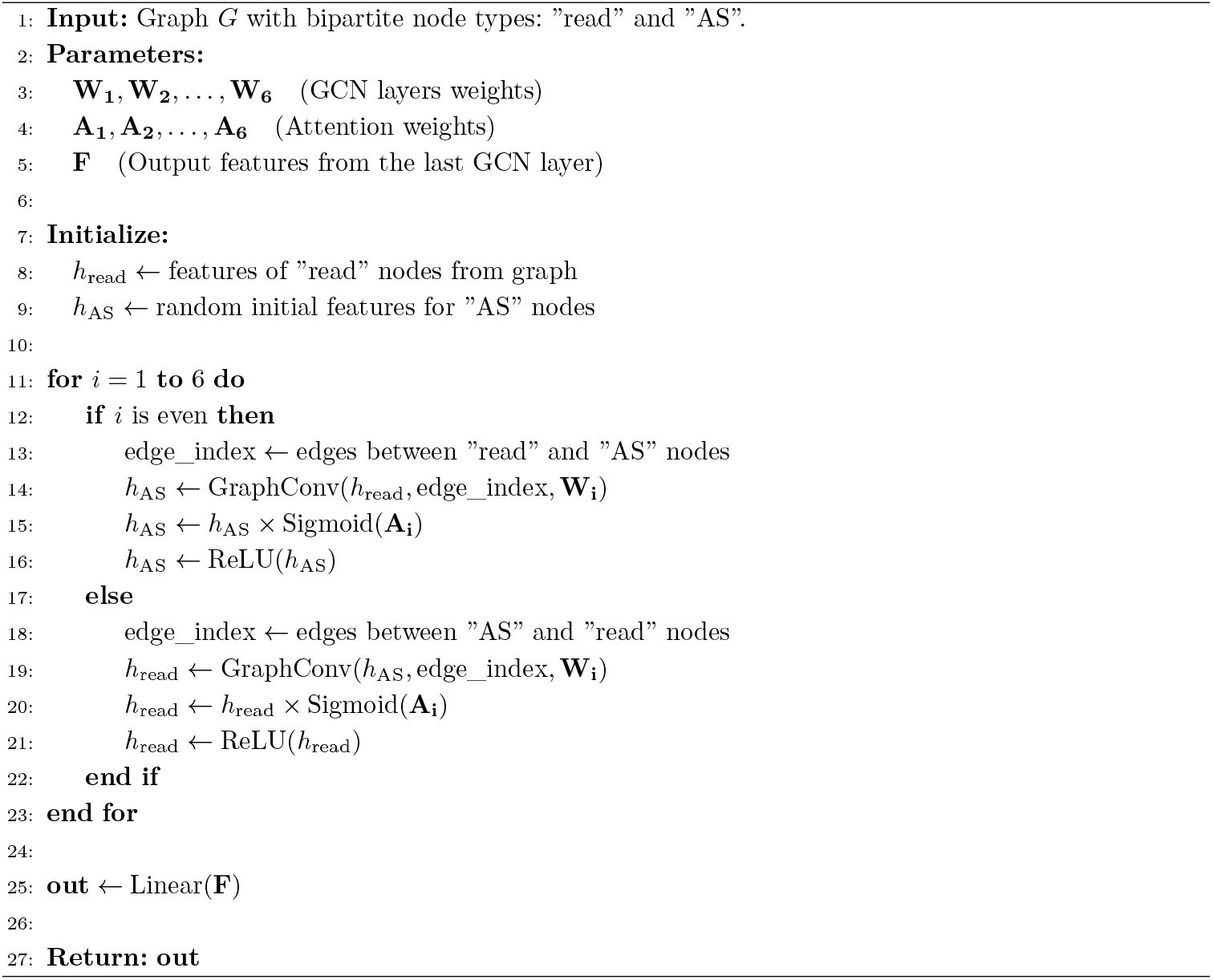

Additionally, the model incorporates a layer-wise attention mechanism. After each graph convolution, the model multiplies the result by a weight and applies the Sigmoid activation function to compute the attention mechanism’s value.

Specifically, the output *h*_*l*_ at layer *l* is computed through graph convolution and the attention mechanism, as given by the following equation:

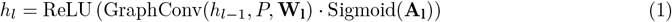

where:

- *h*_*l*−1_ is the output from the previous layer, initially set as the input features.
- *P* is the adjacency matrix representing the graph structure.
- **W**_**l**_ is the attention weight matrix for layer *l*.
- Sigmoid(**A**_**l**_) is the activation of the attention weights, which adjusts the contribution of that layer. The pseudocode **Algorithm 1** provides a detailed introduction to the Bipartite Graph GCN Model.

### 3.5 Validation set and Ensemble Model

We set up a validation set during the training process. At the end of every 10 epochs, the model’s performance was evaluated by calculating the F1-score of the validation set, and the model corresponding to the epoch with the highest F1-score was saved. During the testing phase, predictions were made using multiple trained models, and their output probabilities were averaged through a weighted contribution based on their performance. Let *n* denote the number of models, where each model has an F1-score *f*_*i*_ on the validation set and an output probability *P*_*i*_. The final predicted probability is calculated as follows:

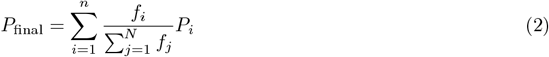

Then, the final predicted label is determined by selecting the class with the highest probability:

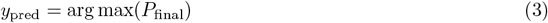

### 3.6 Alignment, assembly, benchmark and evaluation

First, we began by aligning the RNA-seq data to the reference genome using Minimap2 to obtain alignment information. In this step, Minimap2 was used the “splice” mode, which was specifically designed for transcriptome data. This mode effectively identified splice sites and ensures proper handling of sequences that spanned exon-intron boundaries. The resulting SAM file contained the alignment information between the RNA-seq data and the reference genome. By default, all RNA-seq data was mapped to the reference genome with the following command: minimap2 -ax splice reference.fa sample.fastq > sample.sam

Next, we processed the SAM file using samtools [31]. The following command was used to sort and convert the SAM file into BAM format: samtools sort -o sample.bam sample.sam Then, transcript assembly was performed using StringTie2 (version 2.0). StringTie2 utilized the alignment data for transcript assembly, quantification, and structural annotation. In this step, the ‘-L’ option enabled the long-read splicing mode. The command used was: stringtie sample.bam -L -o sample.gtf

Additionally, we conducted a benchmark test using the EASTR software with the following command: eastr –bam sample.bam -r reference.fa -i bowtie2_index_path –out_filtered_bam

To quantitatively assessed the reference intron and exon ratio in the StringTie2 assembly and further verified matching intron chains accuracy, we used the gffcompare to compare the StringTie2 GTF file with the reference gene annotation file. The following command was used: gffcompare -r annotation.gtf -o sample_result sample.gtf

## 4 Data and materials

The experimental data utilized in this study were sourced from publicly available NCBI datasets. Both the training and validation sets consisted of data from Homo sapiens. The test sets included datasets from Homo sapiens, Mus musculus, and Arabidopsis thaliana. All data and genome-related files are accessible for download from the NCBI website (https://www.ncbi.nlm.nih.gov/), with dataset identifiers provided in **Table 2**.

The source code for the latest version of the GLASS package, along with the statistical and assembly results from each tool and the benchmarking pipeline described in this study, are freely available at GitHub: https://github.com/sduljh/GLASS.

## 5 Discussion

In this study, we presented GLASS, a innovative method based on graph model for efficiently filtering alignment files in third-generation RNA-seq. GLASS constructed the ‘Read-AS Map’ model, which allowed for more accurate identification and removal of low-quality alignments, thereby improving the overall quality of the AS information. Our experimental results demonstrated that GLASS had significant advantages when handling complex transcriptome data. Compared to existing filtering tools, it better preserved high-quality transcript information while effectively eliminating noise and redundant data.

Through training on Homo sapiens datasets, this study has also shown good generalization performance in other species such as Mus musculus and Arabidopsis thaliana. The research has the potential to identify errors in alignments for species without reference annotations, offering more accurate and precise transcriptome assembly results for these species. This could help advance research into the transcriptional mechanisms of non-reference annotated organisms.

In recent years, with the rapid advancement of deep learning, significant applications have been achieved in bioinformatics area [32]. However, models based deep learning for third generation RNA-seq are still in their early stages and have not yet fully realized their potential. This study, as an important attempt to utilize deep learning for RNA-seq, not only promises to improve the accuracy of RNA alignments to reference genomes but also holds the potential to provide a framework and reference for future research in more bioinformatics fields, further promoting the application of artificial intelligence in life sciences.

Furthermore, GLASS is not limited to quality control in third-generation transcriptome data; it has great potential for expansion and can be adapted to other domains.

In recent years, single-cell transcriptomics and spatial transcriptomics have seen significant development and are widely applied in cutting-edge biomedical research [33]. In both of these technologies, genome alignment remains a crucial step in data processing. Current single-cell data processing pipelines typically employ two types of filtering techniques: traditional filtering software (e.g., fastp) and filtering based on cells or genes after generating gene expression matrices [34]. However, filtering tools based on alignment files are relatively scarce, and there has been limited discussion in the literature on this topic. Therefore, developing alignment file-based filtering tools is expected to fill this technological gap and provide a more precise and efficient solution for the quantitative analysis of single-cell and spatial transcriptomics.

Moreover, the method proposed in this paper for classifying reads represents a bold attempt. In recent years, long-read sequencing has made some progress in other fields, such as the development of tools like Strainy [35], which has contributed to advances in metagenomic assembly. This brings forth an important and challenging question: how can we further generalize our designed method to enhance the applicability of read classification.

Although the GNN-based algorithm in this study performed excellently in filtering third-generation transcriptome data, there was still room for further optimization. For instance, we can enhance the graph structure introduce by incorporating more biological prior knowledge, design more effective features by introducing sequence information, and apply more complex model architectures to improve the learning performance of the network. Future research also can focus on how to leverage the interpretability of the GNN model to further increase its application value in biological research.

## 6 Conclusion

The innovation of this study lay in the development of a new method for filtering transcriptome alignment files, specifically designed for the characteristics of third-generation RNA-Seq data. By leveraging the long-read nature of third-generation RNA-seq and combining graph machine learning, we have purified the alignment files. Experimental results showed that this approach effectively reduced the rate of non-reference AS and improved the accuracy of transcript assembly of third-generation RNA-Seq. Our work provides a novel perspective for third-generation transcriptomics analysis and offers technical support for future data analysis and tool development in transcriptomics research.

## Supplementary information

Supplementary data are available.

## Author contribution

T.Y., G.L., and J.L. conceived and designed the experiments. J.L., S.G., and Z.G. performed the experiments. T.Y., J.L., and S.G. analyzed the data. J.L., T.Y., J.M., S.G., and G.L. contributed reagents/materials/analysis tools. T.Y., J.L., and G.L. wrote the paper. T.Y. and J.L. designed the software used in the analysis. G.L. oversaw the project.

## References

[1] Marx, V.: Method of the year: long-read sequencing. Nature Methods 20(1), 6–11 (2023)

[2] Kovaka, S., Ou, S., Jenike, K.M., Schatz, M.C.: Approaching complete genomes, transcriptomes and epi-omes with accurate long-read sequencing. Nature methods 20(1), 12–16 (2023)

[3] Conesa, A., Madrigal, P., Tarazona, S., Gomez-Cabrero, D., Cervera, A., McPherson, A., Szczesniak, M.W., Gaffney, D.J., Elo, L.L., Zhang, X., et al.: A survey of best practices for rna-seq data analysis. Genome biology 17, 1–19 (2016)

[4] Stark, R., Grzelak, M., Hadfield, J.: Rna sequencing: the teenage years. Nature Reviews Genetics 20(11), 631–656 (2019)

[5] Marguerat, S., Bähler, J.: Rna-seq: from technology to biology. Cellular and molecular life sciences 67, 569–579 (2010)

[6] Al’Khafaji, A.M., Smith, J.T., Garimella, K.V., Babadi, M., Popic, V., Sade-Feldman, M., Gatzen, M., Sarkizova, S., Schwartz, M.A., Blaum, E.M., et al.: High-throughput rna isoform sequencing using programmed cdna concatenation. Nature Biotechnology 42(4), 582–586 (2024)

[7] Wang, Y., Zhao, Y., Bollas, A., Wang, Y., Au, K.F.: Nanopore sequencing technology, bioinformatics and applications. Nature biotechnology 39(11), 1348–1365 (2021)

[8] Uapinyoying, P., Goecks, J., Knoblach, S.M., Panchapakesan, K., Bonnemann, C.G., Partridge, T.A., Jaiswal, J.K., Hoffman, E.P.: A long-read rna-seq approach to identify novel transcripts of very large genes. Genome research 30(6), 885–897 (2020)

[9] Kim, D., Paggi, J.M., Park, C., Bennett, C., Salzberg, S.L.: Graph-based genome alignment and genotyping with hisat2 and hisat-genotype. Nature biotechnology 37(8), 907–915 (2019)

[10] Dobin, A., Davis, C.A., Schlesinger, F., Drenkow, J., Zaleski, C., Jha, S., Batut, P., Chaisson, M., Gingeras, T.R.: Star: ultrafast universal rna-seq aligner. Bioinformatics 29(1), 15–21 (2013)

[11] Li, H.: Minimap2: pairwise alignment for nucleotide sequences. Bioinformatics 34(18), 3094–3100 (2018)

[12] Han, J., Xiong, J., Wang, D., Fu, X.-D.: Pre-mrna splicing: where and when in the nucleus. Trends in cell biology 21(6), 336–343 (2011)

[13] Shinder, I., Hu, R., Ji, H.J., Chao, K.-H., Pertea, M.: Eastr: Identifying and eliminating systematic alignment errors in multi-exon genes. Nature communications 14(1), 7223 (2023)

[14] Jaganathan, K., Panagiotopoulou, S.K., McRae, J.F., Darbandi, S.F., Knowles, D., Li, Y.I., Kosmicki, J.A., Arbelaez, J., Cui, W., Schwartz, G.B., et al.: Predicting splicing from primary sequence with deep learning. Cell 176(3), 535–548 (2019)

[15] Chao, K.-H., Mao, A., Salzberg, S.L., Pertea, M.: Splam: a deep-learning-based splice site predictor that improves spliced alignments. Genome biology 25(1), 243 (2024)

[16] Xu, C., Bao, S., Wang, Y., Li, W., Chen, H., Shen, Y., Jiang, T., Zhang, C.: Reference-informed prediction of alternative splicing and splicing-altering mutations from sequences. Genome Research 34(7), 1052–1065 (2024)

[17] Smalter, A., Huan, J., Lushington, G.: Graph wavelet alignment kernels for drug virtual screening. Journal of Bioinformatics and computational biology 7(03), 473–497 (2009)

[18] Mahé, P., Vert, J.-P.: Graph kernels based on tree patterns for molecules. Machine learning 75(1), 3–35 (2009)

[19] Hosmer Jr, D.W., Lemeshow, S., Sturdivant, R.X.: Applied Logistic Regression, 3rd edn. Wiley Series in Probability and Statistics. John Wiley & Sons, Hoboken, NJ (2013)

[20] Webb, G.I., Keogh, E., Miikkulainen, R.: Naïve bayes. Encyclopedia of machine learning 15(1), 713–714 (2010)

[21] Breiman, L.: Random forests. Machine learning 45, 5–32 (2001)

[22] Chen, T., Guestrin, C.: Xgboost: A scalable tree boosting system. In: Proceedings of the 22nd Acm Sigkdd International Conference on Knowledge Discovery and Data Mining, pp. 785–794 (2016)

[23] Gurney, K.: An Introduction to Neural Networks, 1st edn. CRC Press Revivals, p. 320. CRC Press, Boca Raton, FL (2018)

[24] Pertea, M., Kim, D., Pertea, G.M., Leek, J.T., Salzberg, S.L.: Transcript-level expression analysis of rna-seq experiments with hisat, stringtie and ballgown. Nature protocols 11(9), 1650–1667 (2016)

[25] Engström, P.G., Steijger, T., Sipos, B., Grant, G.R., Kahles, A., Rätsch, G., Goldman, N., Hubbard, T.J., Harrow, J., et al.: Systematic evaluation of spliced alignment programs for rna-seq data. Nature methods 10(12), 1185–1191 (2013)

[26] Kovaka, S., Zimin, A.V., Pertea, G.M., Razaghi, R., Salzberg, S.L., Pertea, M.: Transcriptome assembly from long-read rna-seq alignments with stringtie2. Genome biology 20, 1–13 (2019)

[27] Pertea, G., Pertea, M.: Gff utilities: Gffread and gffcompare. F1000Research 9 (2020)

[28] Kipf, T.N., Welling, M.: Semi-supervised classification with graph convolutional networks. arXiv preprint arXiv:1609.02907 (2016)

[29] Zhang, W., Yin, Z., Sheng, Z., Li, Y., Ouyang, W., Li, X., Tao, Y., Yang, Z., Cui, B.: Graph attention multi-layer perceptron. In: Proceedings of the 28th ACM SIGKDD Conference on Knowledge Discovery and Data Mining, pp. 4560–4570 (2022)

[30] Müller, R., Kornblith, S., Hinton, G.E.: When does label smoothing help? Advances in neural information processing systems 32 (2019)

[31] Li, H., Handsaker, B., Wysoker, A., Fennell, T., Ruan, J., Homer, N., Marth, G., Abecasis, G., Durbin, R., Subgroup,. G.P.D.P.: The sequence alignment/map format and samtools. bioinformatics 25(16), 2078–2079 (2009)

[32] Alam, S., Israr, J., Kumar, A.: In: Singh, V., Kumar, A. (eds.) Artificial Intelligence and Machine Learning in Bioinformatics, pp. 321–345. Springer, Singapore (2024)

[33] Gulati, G.S., D’Silva, J.P., Liu, Y., Wang, L., Newman, A.M.: Profiling cell identity and tissue architecture with single-cell and spatial transcriptomics. Nature Reviews Molecular Cell Biology, 1–21 (2024)

[34] Butler, A., Hoffman, P., Smibert, P., Papalexi, E., Satija, R.: Integrating single-cell transcriptomic data across different conditions, technologies, and species. Nature biotechnology 36(5), 411–420 (2018)

[35] Kazantseva, E., Donmez, A., Frolova, M., Pop, M., Kolmogorov, M.: Strainy: phasing and assembly of strain haplotypes from long-read metagenome sequencing. Nature Methods 21(11), 2034–2043 (2024)

